# Prophylactic inhibition of colonization by *Streptococcus pneumoniae* with the secondary bile acid metabolite deoxycholic acid

**DOI:** 10.1101/2021.05.17.444594

**Authors:** Jorge E. Vidal, Meagan N. Wier, Uriel Angulo-Zamudio, Erin McDevitt, Ana G. Jop Vidal, Babek Alibayov, Anna Scasny, Sandy M. Wong, Brian J. Akerley, Larry S. McDaniel

## Abstract

Streptococcus *pneumoniae* (Spn) colonizes the nasopharynx of children and the elderly but also kills millions worldwide yearly. The secondary bile acid metabolite, deoxycholic acid (DoC), affects the viability of human pathogens but also plays multiple roles in host physiology. We assessed *in vitro* the antimicrobial activity of DoC and investigated its potential to eradicate Spn colonization using an *ex vivo* model of human nasopharyngeal colonization and an *in vivo* mouse model of colonization. At a physiological concentration DoC (0.5 mg/ml; 1.27 mM) killed all tested Spn strains (N=48) two h post-inoculation. The *ex-vivo* model of nasopharyngeal colonization showed that DoC eradicated colonization by Spn strains as soon as 10 min post-exposure. The mechanism of action did not involve activation of autolysis since the autolysis-defective double mutants Δ*lytA*Δ*lytC* and *ΔspxBΔlctO* were as susceptible to DoC as was the wild-type (WT). Oral streptococcal species (N=20), however, were not susceptible to DoC (0.5 mg/ml). Unlike trimethoprim, whose spontaneous resistance frequency (srF) for TIGR4 or EF3030 was ≥1×10^−9^, no spontaneous resistance was observed with DoC (srF≥1×10^−12^). Finally, the efficacy of DoC to eradicate Spn colonization was assessed *in vivo* using a topical route via intranasal (i.n.) administration and as a prophylactic treatment. Mice challenged with Spn EF3030 carried a median of 4.05×10^5^ cfu/ml four days post-inoculation compared to 6.67×10^4^ cfu/ml for mice treated with DoC. Mice in the prophylactic group had a ∼99% reduction of the pneumococcal density (median, 2.61 ×10^3^ cfu/ml). Thus, DoC, an endogenous human bile salt, has therapeutic potential against Spn.

## Introduction

*Streptococcus pneumoniae* (Spn) colonizes millions worldwide yearly, with a colonization prevalence particularly high in children and the elderly (1-3). In children, the prevalence of pneumococcal carriage can be as high as 90% in those from developing countries, or between 25-40% in children from industrialized nations (1, 4-7). There are also certain populations within developed countries with increased carriage rates. For example, in a study by Sutcliffe et al (2019) 73.5% of children less than five, living in the US but of the Navajo nation, carried the pneumococcus in the upper airways (8). Along the same lines, carriage of Spn in adults 18-49.9 years of age can be as high as 50% (8, 9) while pneumococcal carriage in a more vulnerable population, those older than 60, is similar (10) but increases in individuals colonized by the influenza virus (5). Although it is early to draw conclusions, pneumococcal carriage is expected to increase in individuals colonized with SARS-CoV-2 whereby we may experience a surge of pneumococcal disease (PD) cases in the next few years (11).

Pneumococcal carriage is a risk for developing PD and therefore it is considered an immediate and necessary precursor of PD (1, 2, 12, 13). An important intervention with a demonstrated positive impact on Spn carriage has been vaccination (14). Pneumococcal conjugated vaccines (PCV) were introduced in many parts of the world since 2001 when PCV7 was licensed in the US (15, 16). The introduction of these vaccines reduced the burden of PD caused by vaccine serotypes on a global scale and has also decreased nasopharyngeal carriage of pneumococcal vaccine types in vaccinated populations (14, 17). However, the overall carriage prevalence has not changed because of a phenomenon called “serotype replacement”, i.e., vaccine-escape strains have replaced vaccine type (VT) strains in the nasopharynx, resulting in pneumococcal carriage rates similar to those observed prior to the introduction of vaccines (18-20). Therefore, additional interventions, or prophylactic strategies, are needed to aid reduce the burden of colonization.

Pathogenic and normal flora bacteria are susceptible to bile from different mammals (21-23). Recent studies have demonstrated that the lack of primary and secondary bile acid metabolites is implicated in the development of intestinal infectious disease (21-23). Bile consists of ∼95% water in which are dissolved a number of endogenous solid constituents including bile salts, bilirubin phospholipid, cholesterol, amino acids, steroids, enzymes, porphyrins, vitamins, and heavy metals (24). Bile salts are the major organic solutes in bile and normally function to emulsify dietary fats and facilitate their intestinal absorption. The two main primary bile salts that are synthesized in the mammalian liver are cholic acid, and chenodeoxycholic acid (24, 25). Intestinal bacteria then produce “secondary bile acids” through enzymatic reactions, with the addition of two hydroxyl groups to cholic acid producing one of the most abundant secondary metabolites, deoxycholic acid (DoC) (24, 26). Endogenously-produced, secondary bile acids, have widespread effects on the host and resident microbiota and have therefore been used to treat different diseases (26). Cholic acid is used to treat patients with genetic deficiencies in the synthesis of bile acids due to single enzyme deficiencies; the typical dose is 10 to 15 mg/kg once daily (27). DoC has been utilized in humans at a concentration of 15 mg/kg/day to decrease plasma high-density lipoprotein (HDL)-cholesterol and low-density lipoprotein (LDL)-cholesterol (27) and it is also FDA-approved to reduce fat deposits (28, 29).

In addition to their physiological role in digesting lipids, bile acids are important regulators of intestinal homeostasis by activating receptors on intestinal cells and stimulating the immune response (25). In germ-free mice or mice treated with antibiotics to deplete the intestinal bacterial flora, the lack of secondary bile acids caused a deficient TLR7-MyD88 signaling in plasmacytoid dendritic cells that resulted in an increased susceptibility to systemic chikungunya infection (21). Another secondary bile acid, ursodeoxycholic acid, inhibited *in vitro* toxin production, growth and spore germination of *Clostridiodes difficile* strains and contributed to colonization resistance in an *in vivo* mouse model of *C. difficile* disease (CDI) (30). Administration of clindamycin reduced intestinal levels of valerate and DoC, increasing viable counts of *C. difficile* in a CDI chemostat infection model (31). A similar treatment of mice with clindamycin caused decreased intestinal DoC leading to increased *Campylobacter jejuni*-induced colitis (22). The growth of another intestinal pathogen, *Clostridium perfringens*, was inhibited *in vitro* with as low as 50 µM of DoC. Using an animal model of necrotic enteritis (NE), and supplementation of DoC in the diet (e.g., 1.5 g/kg) decreased intestinal inflammation and NE-associated intestinal cell death and apoptosis (23).

*In vitro* studies demonstrated synergistic antibiotic activity when Doc (2.5 mg/ml) was combined with vancomycin, or with vancomycin and furazolidone, to treat clinically-important antibiotic resistant pathogens (32). Synergism between DoC, or lithocholic acid, and tryptophan-derived antibiotics secreted by intestinal bacteria inhibited bacterial division of *C. difficile* strains (33). In this study we assessed the antimicrobial activity of DoC against a collection of Spn strains, including reference strains, Spn isolated from cases of pneumococcal disease, and strains resistant to multiple antibiotics. The mechanistic basis for the observed sensitivity to DoC was investigated using knockout mutants in different autolytic mechanisms. We also adapted an *ex vivo* model of nasopharyngeal colonization to investigate the efficacy of DoC to eradicate Spn colonization and finally developed an *in vivo* prophylactic mouse model to demonstrate that administration of DoC prevented Spn colonization.

## Results

### Deoxycholic acid (DoC) eradicates cultures of drug-resistant *S. pneumoniae*

We challenged reference strains D39 and TIGR4 and drug-resistant Spn 19A and 19F vaccine-serotypes strains with increasing dosages of DoC and incubated for 2 h. Whereas untreated bacteria grew at >10^8^ cfu/ml, all five pneumococcal strains treated with DoC were killed within 2 h with 0.5 mg/ml (1.27 mM). Except for strain D39, all other vaccine serotype Spn strains were also killed with 250 µg/ml of DoC (Fig. 1).

**Figure 1.**
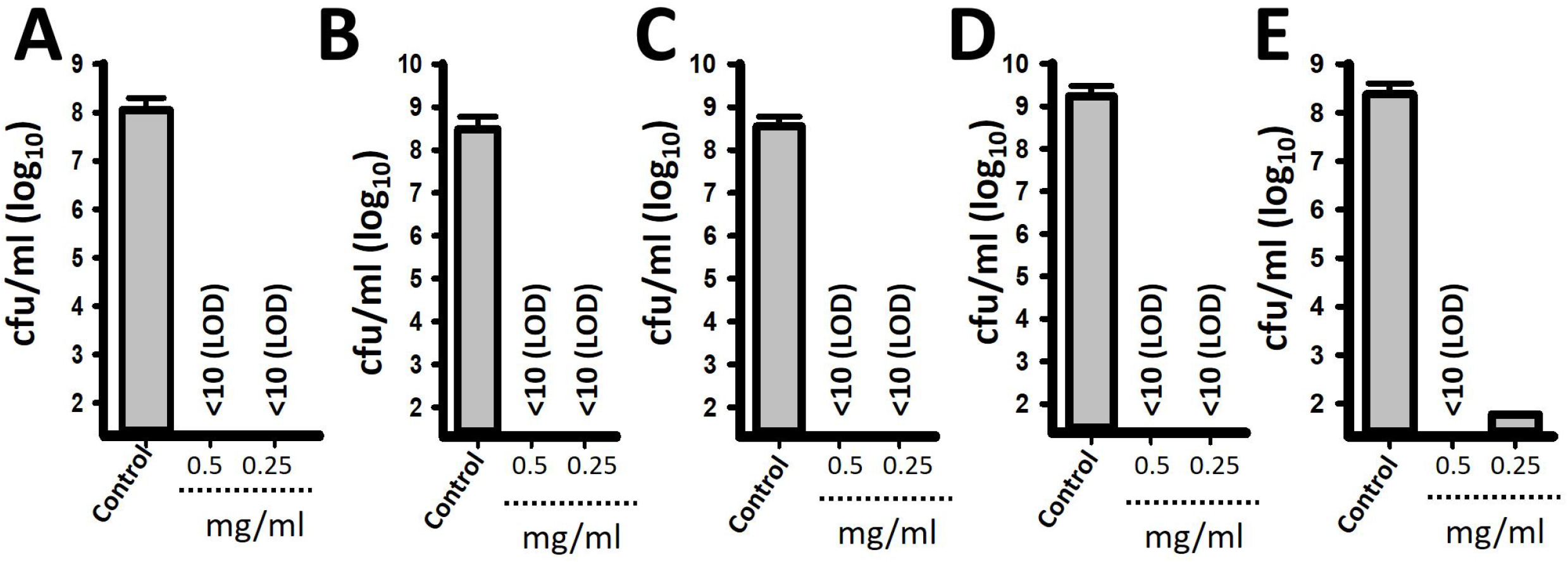
Deoxycholic acid kills *S. pneumoniae* strains within two hours of incubation. *S. pneumoniae* strain (A) GA47281, (B) GA44288, (C) GA17227, (D) TIGR4, or (E) D39 was inoculated at a density of ∼5.15×10^8^ cfu/ml in THY broth and left untreated (control) or treated with 0.5 mg/ml or 0.25 mg/ml of deoxycholic acid (DoC). Bacteria were incubated for 2 h at 37°C in a 5% CO_2_ atmosphere after which the cultures were serially diluted and plated onto blood agar plates to obtain the density (cfu/ml). Error bars represent the standard errors of the means calculated using data from at least three independent experiments. The limit of detection (LOD) was <10 cfu/ml.

We then assess the antimicrobial efficacy of DoC (0.5 mg/ml) against 39 pneumococcal strains isolated from cases of pneumococcal disease. Strains represented all PCV13 vaccine serotypes, and at least two strains of each vaccine serotype were challenged. Supplemental Table 1 confirmed that 0.5 mg/ml (1.27 mM) of DoC incubated for 2 h killed all assessed Spn strains. A MIC_90_ of 0.5 mg/ml DoC was established.

A similar MIC [0.5 mg/ml (1.27 mM)] was obtained with reference strain TIGR4, or EF3030, assessed in cation-activated Mueller-Hinton broth (CAMHB) with 3% of lysed horse blood (LHB) and according to CLSI guidelines to assess resistance (34), or non-susceptibility, of Spn strains (Supplemental Fig. 1 and not shown). Whereas the density of TIGR4 inoculated in CAMHB and incubated overnight reached a density of 6.7×10^10^ cfu/ml, those cultures treated with DoC (1.27 mM) had a median density of 1.2×10^4^ cfu/ml, a ∼99.99% reduction in density after overnight incubation (Supplemental Fig. 1).

### DoC eradicates MDR *S. pneumoniae* colonization in an *ex-vivo* model with human pharyngeal cells

To assess the potential of DoC to eradicate pneumococcal colonization, we performed experiments using an *ex-vivo* model of colonization. To create an abiotic substrate for pneumococci to colonize we used polarized human pharyngeal cells that had been fixed with paraformaldehyde. These cell monolayers were infected with a MDR Spn strain GA47281, a vaccine strain serotype 19F bearing resistance to several antibiotics including erythromycin, meropenem, and cefuroxime. Infected human cells were incubated for 4 h to allow attachment of Spn and then planktonic bacteria were removed. Spn-colonized human cells were left untreated or challenged with different dosages of DoC and incubated for 2 h. All doses of DoC reduced the density of MDR pneumococci >90% after the 2 h incubation compared to untreated human pharyngeal cells which had a density of ∼1×10^7^ cfu/ml of pneumococci (Fig. 2A). GA47281 bacteria were not recovered when pneumococci were treated with 0.5 mg/ml (1.27 mM). Confocal scanning laser microscopy, XY optical sections and 3D reconstructions, of cells infected with Spn confirmed that DoC had eradicated colonization by strain GA47281 (Fig. 2C) or colonization by strain TIGR4 (Fig. 2D) from human pharyngeal cells.

**Figure 2.**
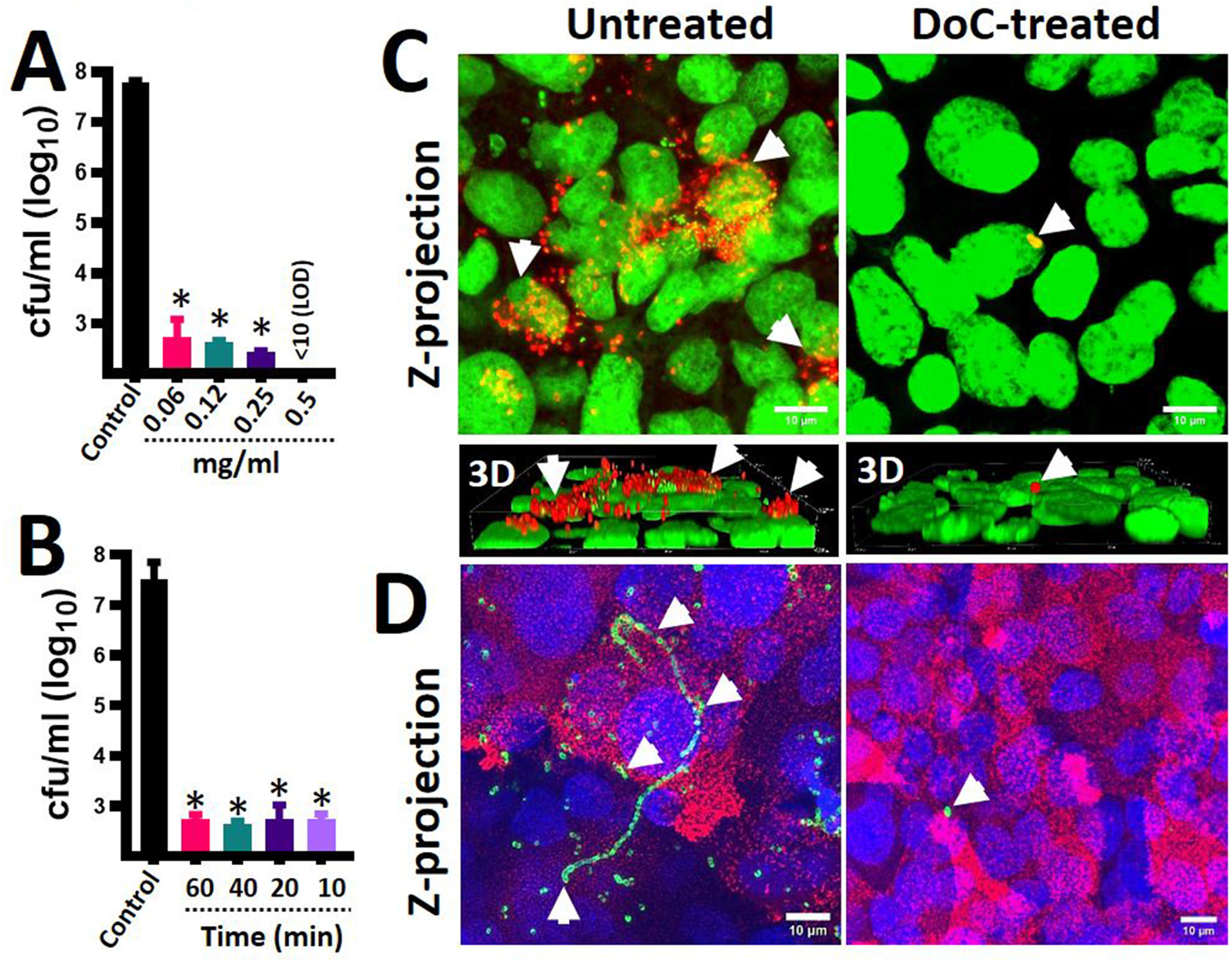
Eradication of pneumococcal colonization with DoC using an *ex vivo* model of human pharyngeal colonization. Polarized human pharyngeal Detroit 562 cells were immobilized with 2% paraformaldehyde and then infected with *S. pneumoniae* strain GA47281 (∼5.15×10^8^ cfu/ml). Infected cells were incubated for four hours and planktonic cells were removed. These *S. pneumoniae*-colonized human pharyngeal cells were left untreated (control) or (A) treated with different dosages of deoxycholic acid (DoC) and incubated for an additional 2 h period, or (B) treated with DoC (0.5 mg/ml) and incubated for the indicated time. At the end of the incubation the cultures were serially diluted and plated onto blood agar plates to obtain the density (cfu/ml). In (A and B) error bars represent the standard errors of the means calculated using data from at least three independent experiments. \**p*<0.05 compared to the untreated control. LOD=limit of detection. (C and D) *S. pneumoniae*-colonized human pharyngeal cells (infected as above) were left untreated or treated with DoC (0.5 mg/ml) and incubated for 2 h. Preparations were fixed and Spn was stained with (C) an anti-S19-Alexa-555 labeled antibody (red) and the DNA was stained with TOPRO3 (green) or with (D) an anti-S4-Alexa-488 labeled antibody (green), cell membranes were labeled with WGA (Red) and DNA with DAPI (blue). Preparations were analyzed by a confocal microscope. Panels show z-projections of z-stacks obtained from xy optical sections. Lower panels in (C) show a 3-D digital reconstruction. The merge of the channels is shown in each panel. Arrows point out extracellular *S. pneumoniae* bacteria.

To determine the exposure time required to kill pneumococci, we performed a time course study treating MDR pneumococci colonizing human pharyngeal cells for up to one h with 0.5 mg/ml (1.27 mM). Interestingly, a ten-minute exposure time was enough to kill most attached MDR pneumococci (Fig. 2B). The antimicrobial effect did not change with a longer exposure time of 60 min. These data indicate that 0.5 mg/ml (1.27 mM) of DoC rapidly eradicates MDR pneumococci that otherwise would have colonized human nasopharyngeal cells at a density ∼10^7^ cfu/ml (Fig. 2B).

### Spontaneous resistance to DoC (500 µg/ml) was not developed by *S. pneumoniae* strains

Spn strains develop spontaneous resistance to some antibiotics at a spontaneous mutation frequency >1×10^−8^ (35). We assessed spontaneous resistance to DoC using antibiotic sensitive strains TIGR4 and EF3030. As expected, spontaneous resistance against trimethoprim developed at a frequency of ≥1.39×10^−9^ when either pneumococcal strain was assessed (Fig. 3). Spontaneous resistance to DoC, however, was not observed in any of the two strains tested even at a population density >10^12^ cfu/ml (Fig. 3).

**Figure 3.**
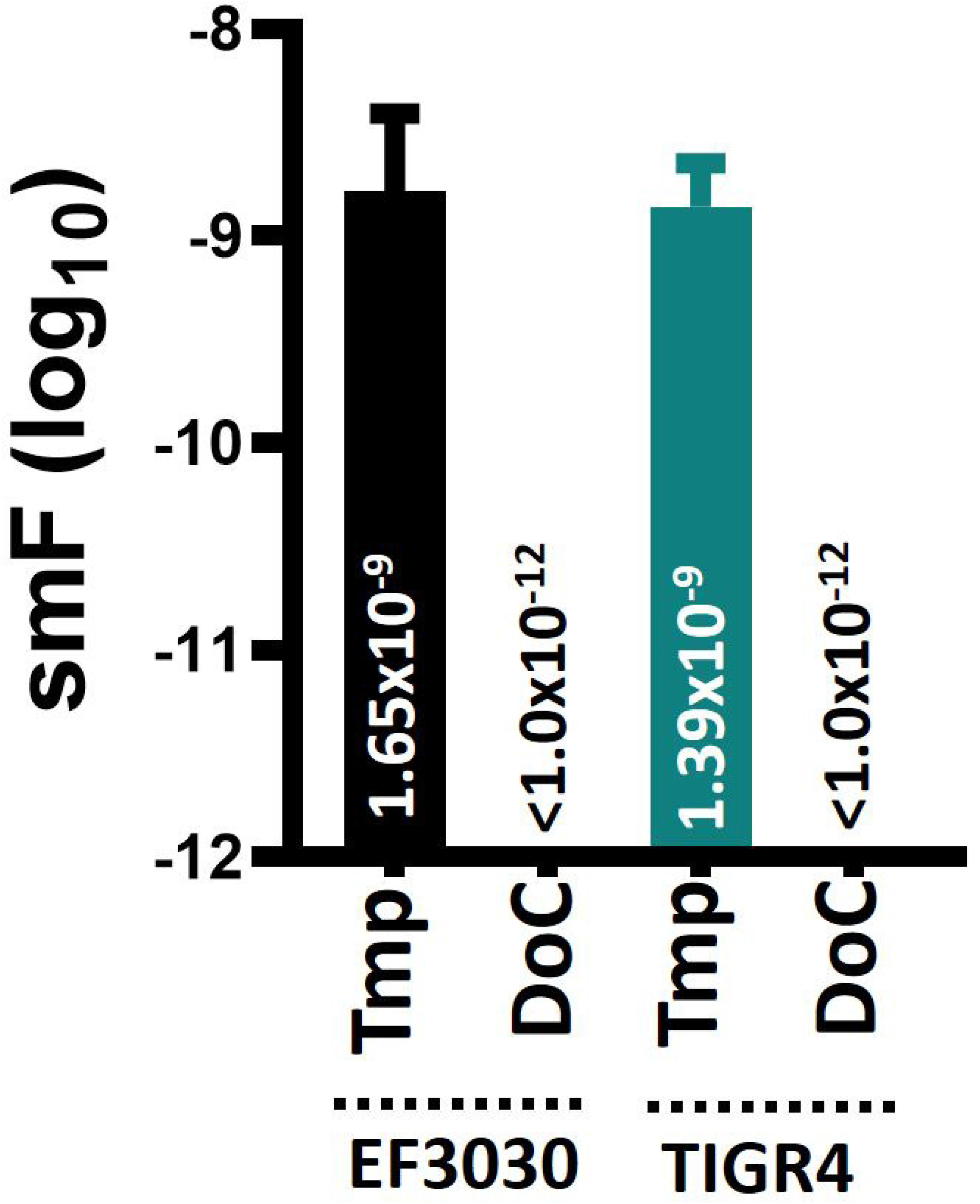
Spontaneous resistance mutation frequency of *S. pneumoniae* for trimethoprim and deoxycholic acid. Bacterial suspensions prepared with fresh cultures of *S. pneumoniae* strain EF3030, or TIGR4, were adjusted to a density of ∼10^8^, ∼10^9^, ∼10^10^, ∼10^11^, and ∼10^12^ cfu/ml and inoculated onto blood agar plates (BAP) containing trimethoprim (Tmp, 1 µg/ml) or deoxycholic acid (DoC, 0.5 mg/ml). Each inoculum was further diluted and plated onto plain BAP. All cultures were incubated for 20 h after which bacteria were counted. The spontaneous mutation frequency (smF) was calculated by dividing the number of spontaneous resistant pneumococci, i.e., grown on BAP with trimethoprim or DoC, by the bacterial population. Error bars represent the standard errors of the means calculated using data from at least three independent experiments. The median smF is shown on each bar.

### The mechanism of DoC-killing does not appear to trigger autolysis

Since the above experiments revealed that a short exposure time was sufficient to kill pneumococci, we hypothesized that an irreversible autolytic mechanism may have been triggered by DoC. Autolysis is mainly driven by autolysins LytA and LytC (36, 37) but hydrogen peroxide produced by the pneumococcus, through enzymes SpxB and LctO, also contributes to lysis of pneumococci (38-40). To assess this hypothesis, we utilized strain R6 wt, its isogenic R6Δ*lytA*Δ*lytC* mutant and a double R6*ΔspxBΔlctO* mutant to assess the role of autolysis in the observed DoC-mediated bactericidal activity. Since autolysis can be measured by quantifying the release of extracellular (e)DNA, we first confirmed that the absence of the LytA and LytC autolysins, or SpxB and LctO, causes a decreased release of eDNA into the supernatant. After four hours of incubation, the eDNA released by R6 wt strain reached a median of 2.24×10^6^ pg/ml, whereas R6*ΔspxBΔlctO* released a statistically different two-fold decreased amount of eDNA (median, 1.09×10^6^ pg/ml) in the supernatant (Fig. 4A). As expected, the autolysin double mutant R6Δ*lytA*Δ*lytC* yielded a ∼14-fold reduced amount of eDNA (median, 1.55×10^5^ pg/ml) (Fig. 4A) compared to R6 wt.

**Figure 4.**
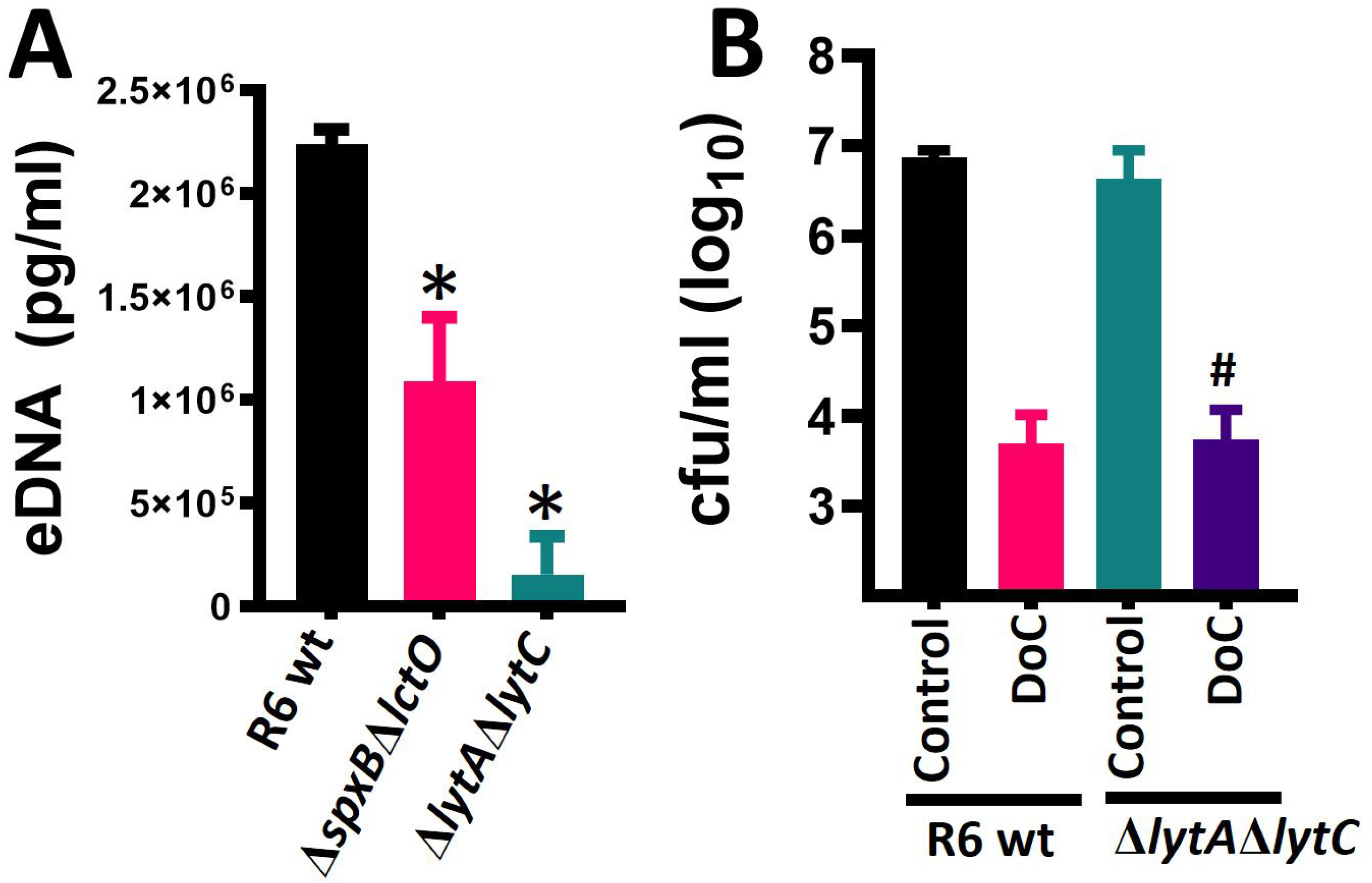
Deoxycholic acid kills pneumococci with a deficient autolytic mechanism. (A) *S. pneumoniae* strain R6 wt, or isogenic double mutants Δ*lytA*Δ*lytC* or Δ*spxB*Δ*lctO* were inoculated in six-well plates containing THY and bacteria were incubated for 4 h. The supernatants were obtained, filter sterilized and eDNA was purified. This eDNA was used as a template in species-specific quantitative (q)PCR reactions along with DNA standards for quantification purposes. *statistical significance, *p*<0.037, compared to R6 wt. (B) R6 wt or its isogenic Δ*lytA*Δ*lytC* mutant was inoculated at a density of ∼5.15×10^8^ cfu/ml in THY broth and left untreated (control) or treated with 0.5 mg/ml of DoC. Bacteria were incubated for 2 h at 37αC in a 5% CO_2_ atmosphere after which the cultures were serially diluted and plated onto blood agar plates to obtain the density (cfu/ml). ^#^*p*=0.33 compared to R6 wt strain treated with DoC. In panels A and B error bars represent the standard errors of the means calculated using data from at least three independent experiments.

These two mutant strains with an impaired autolysis phenotype were then challenged with DoC [0.5 mg/ml (1.27 mM)] and treated and untreated bacteria were incubated for 1 h. Results in Fig. 4B showed a >90% significant reduction (i.e., killing) of the population of R6 pneumococci after 1 h of incubation. The density of the R6*ΔspxBΔlctO* isogenic mutant (not shown) or the R6Δ*lytA*Δ*lytC* mutant (Fig. 4B) was similarly reduced after 1 h incubation period with DoC, indicating that the mechanism by which DoC kill pneumococci is not by triggering autolysis.

### DoC does not affect viability of normal flora streptococci

We next investigated whether the MIC_90_ of DoC that eradicates Spn strains [0.5 mg/ml (1.27 mM)] within 2 h of incubation would have the same bactericidal effect against other streptococci that reside in the oral cavity. As a positive control we utilized Spn reference strain EF3030 (41, 42). The median density of EF3030 untreated cultures was 3.37×10^7^ cfu/ml whereas those treated with 0.5 mg/ml DoC for 2 h had a median density of 3.1×10^2^ cfu/ml, and therefore DoC killed 99.99% of the bacterial population (Fig. 5A). However, the same dose of DoC incubated for 2 h did not significantly affect the viability of *S. oralis* (Fig. 5B), *S. mutans*, (Fig. 5C) *S. gordonii*, (Fig. 5D) and induced a two-log reduction of the density of *S. anginosus*. (Fig. 5E). An additional 16 different streptococcal species were treated with DoC [0.5 mg/ml (1.27 mM)] for two hours and, except for *S. pseudopneumoniae* and *S. salivarius* that were susceptible, cultures of all other species were not affected (Table 1).

**Figure 5.**
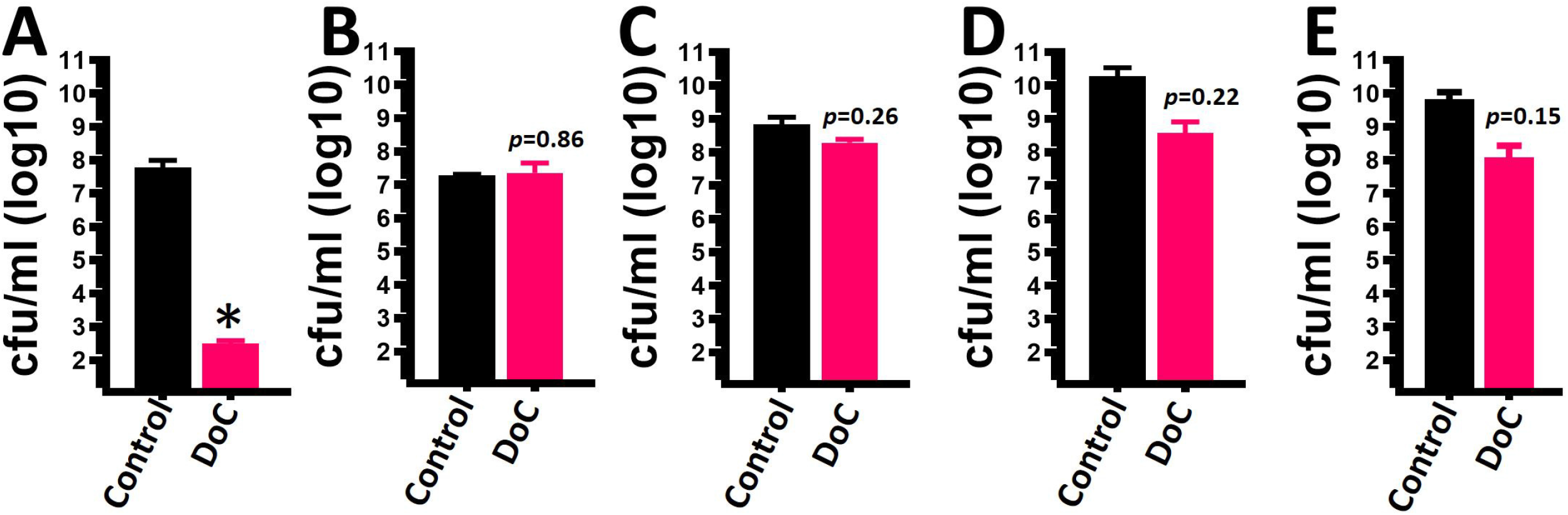
Deoxycholic acid (0.5 mg/ml) does not affect the viability of other streptococcal species. (A) *S. pneumoniae* strain EF3030, (B) *S. oralis*, (C) *S. mutans*, (D) *S. gordonii*, or (E) *S. anginosus* was inoculated at a density of ∼5.15×10^8^ cfu/ml in THY broth and left untreated (control) or treated with 0.5 mg/ml deoxycholic acid (DoC). Bacteria were incubated for 2 h at 37αC in a 5% CO_2_ atmosphere after which the cultures were serially diluted and plated onto blood agar plates to obtain the density (cfu/ml). Error bars represent the standard errors of the means calculated using data from at least three independent experiments. *statistical significance, *p*<0.04, compared to the untreated EF3030 control.

**Table 1.**
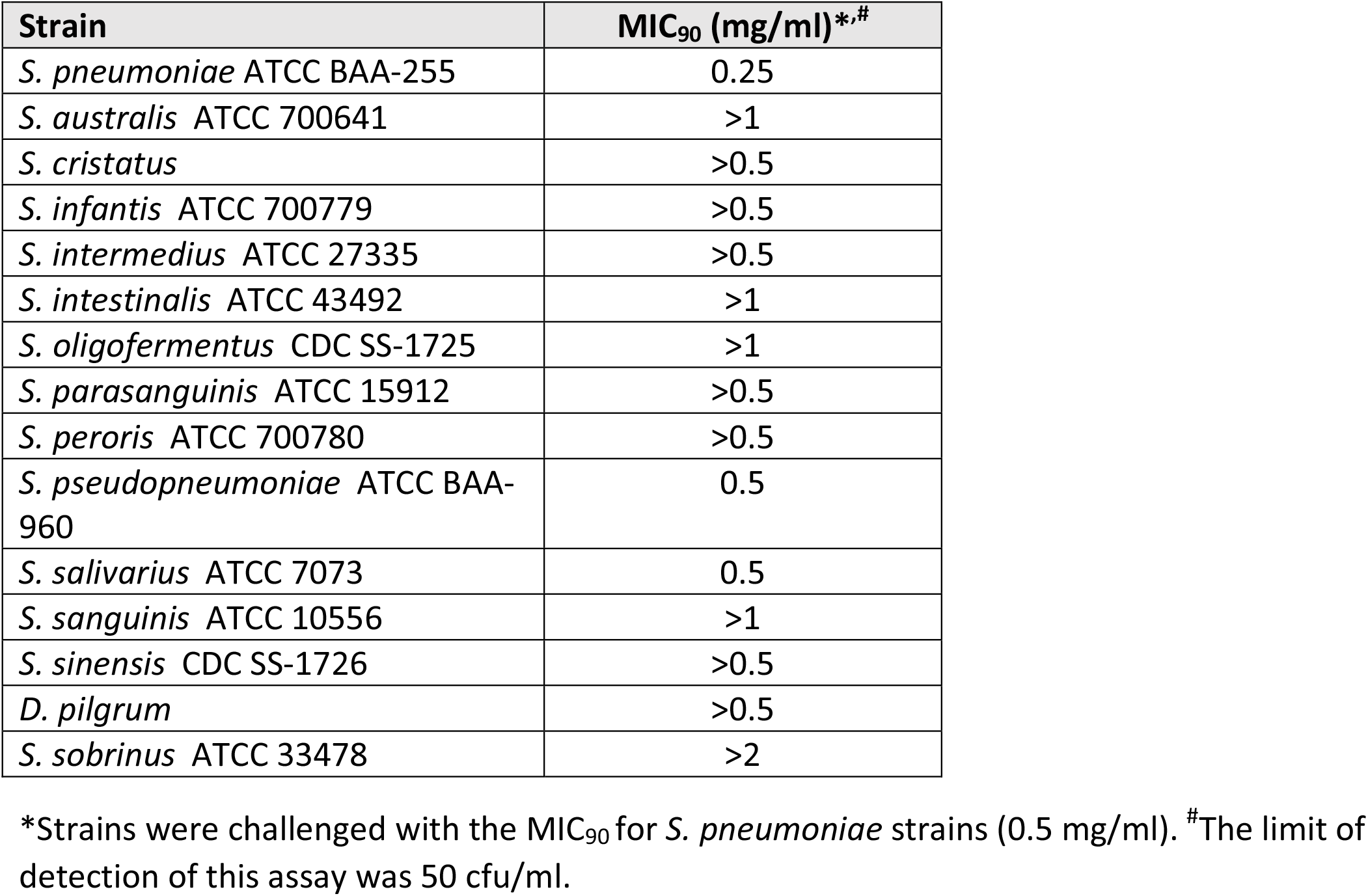
Antimicrobial activity of DoC against streptococcal species.

### Standardizing the mouse model of pneumococcal carriage to assess the efficacy of DoC to eradicate nasopharyngeal colonization

The bacterial inoculum utilized in a mouse model of pneumococcal colonization is usually ∼1×10^7^ cfu and nasal washes through the trachea are performed to assess colonization (43, 44). We first standardized the removal of a defined section of nasopharyngeal tissue consisting of the intact nasal septum (Fig. 6A) from colonized mice and confirmed by histological analysis the presence of typical nasopharyngeal tissue including microvilli, and pseudostratified, squamous epithelium overlaying by loose connective tissue including blood vessels with erythrocytes (Fig. 6B). Since our goal was to assess the efficacy of DoC to reduce, and/or inhibits, nasopharyngeal colonization we investigated a low inoculum density of Spn EF3030 that would sustain colonization of mice, but avoided using a non-natural (i.e., heavy) inoculum. Our experiments demonstrated that inoculating ∼1×10^5^ cfu in the nostrils of mice allowed nasopharyngeal colonization for up to four days at a median density of 2.49×10^5^ cfu/organ (Fig. 6D). We additionally removed the trachea and lungs and demonstrated consistent Spn colonization of the trachea, at a low density of 9.0×10^2^ cfu/organ, but colonization of the lungs was not observed (Fig. 6D). Encapsulated Spn were detected using an anti-S19-Alexa-555 antibody and were identified both in nasopharyngeal homogenates (Fig. 6C) and colonizing the nasopharyngeal tissue (Fig. 6E). Microinvasion into the nasopharyngeal epithelium was also observed (Fig. 6E).

**Figure 6.**
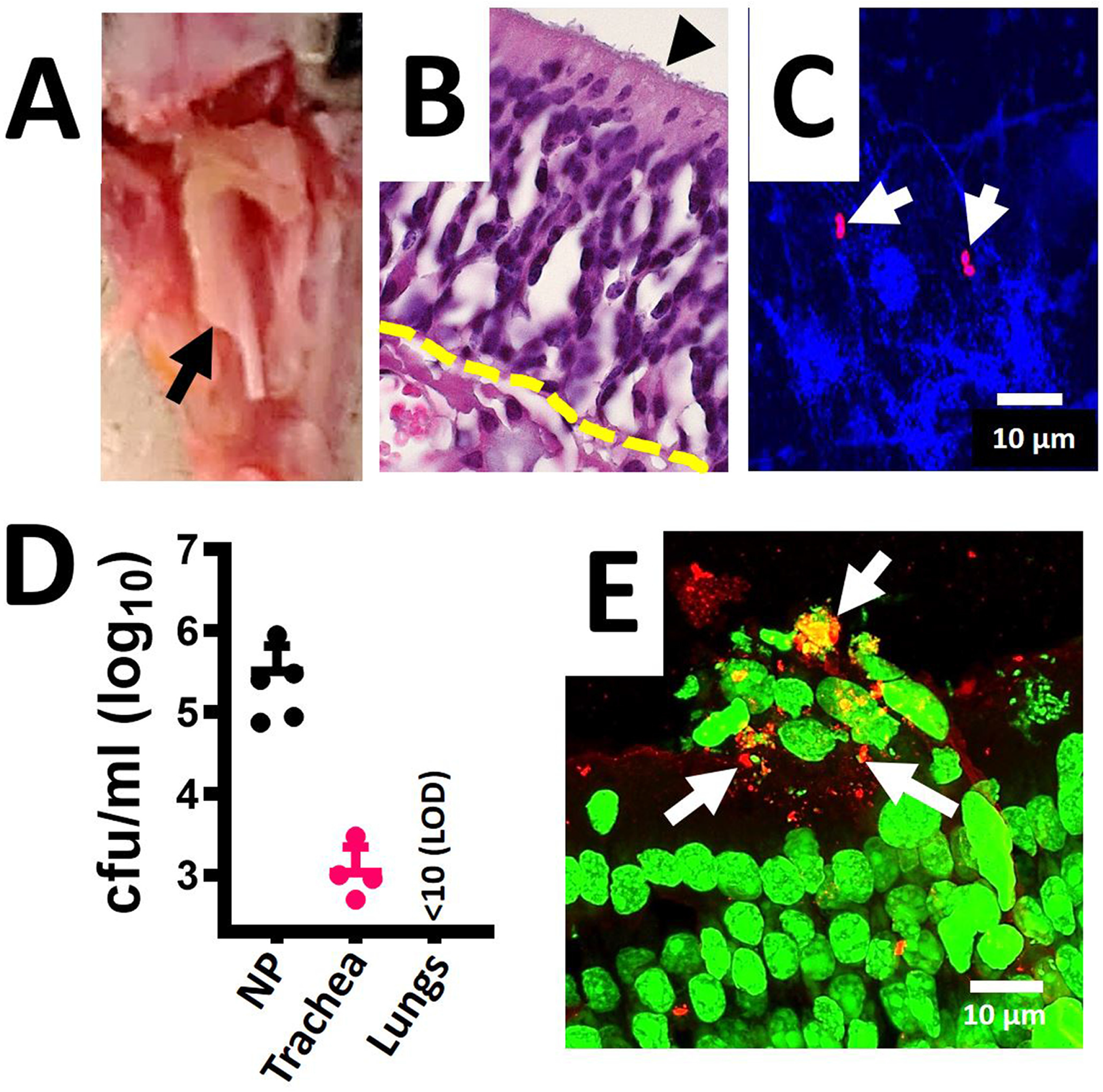
Revisiting the mouse model of pneumococcal nasopharyngeal carriage. C57BL/6 mice (N=5) were intranasally inoculated with *S. pneumoniae* EF3030 (∼1×10^5^ cfu). After 48 h mice were euthanized and the nasal bone was removed to expose the (A) nasopharynx, arrow. The nasopharynx, trachea and lungs were removed. (B) The nasopharynges were sectioned (∼5 µm) and stained with hematoxylin and eosin. Arrowhead=microvilli, dotted line=connective tissue. Nasopharyngeal (NP) tissue, trachea, and lungs were homogenized and (C) NP homogenate was stained with DAPI and with an anti-S19-Alexa-555 antibody, or (D) homogenates were diluted and plated onto BAP with gentamicin (25 µg/ml) to obtain the bacterial density (cfu/ml). (E) NP tissue stained with TOTO-1 and with an anti-S19-Alexa-555 antibody; arrows=pneumococci (red). The micrographs in C and D are z-projections of z-stacks obtained from xy optical sections collected with a confocal microscope.

### Deoxycholic acid (DoC) decreased *in vivo* colonization by *S. pneumoniae* strain EF3030 in a mouse model of nasopharyngeal carriage

We then assessed the efficacy of DoC to eradicate colonization using three groups of mice (N=8). Groups 1 and 2 drank regular water while the drinking water of group 3 was supplemented with DoC at a concentration of 0.2 µg/ml (i.e., 0.02%) six days prior to infection and remained in their drinking water throughout the experiment (Fig. 7A). All three groups were then challenged i.n. with Spn EF3030 (∼1×10^5^ cfu) and 24 h post-infection, mice in groups 1 and 2 were treated twice a day i.n. with 10 ul of a PBS solution or DoC (2 mg/ml), respectively. Mice were euthanized 10 days after initiating the prophylactic regimen (i.e., oral administration) in drinking water, or four days after placebo or DoC topical nasopharyngeal treatment began (Fig. 7A).

**Figure 7.**
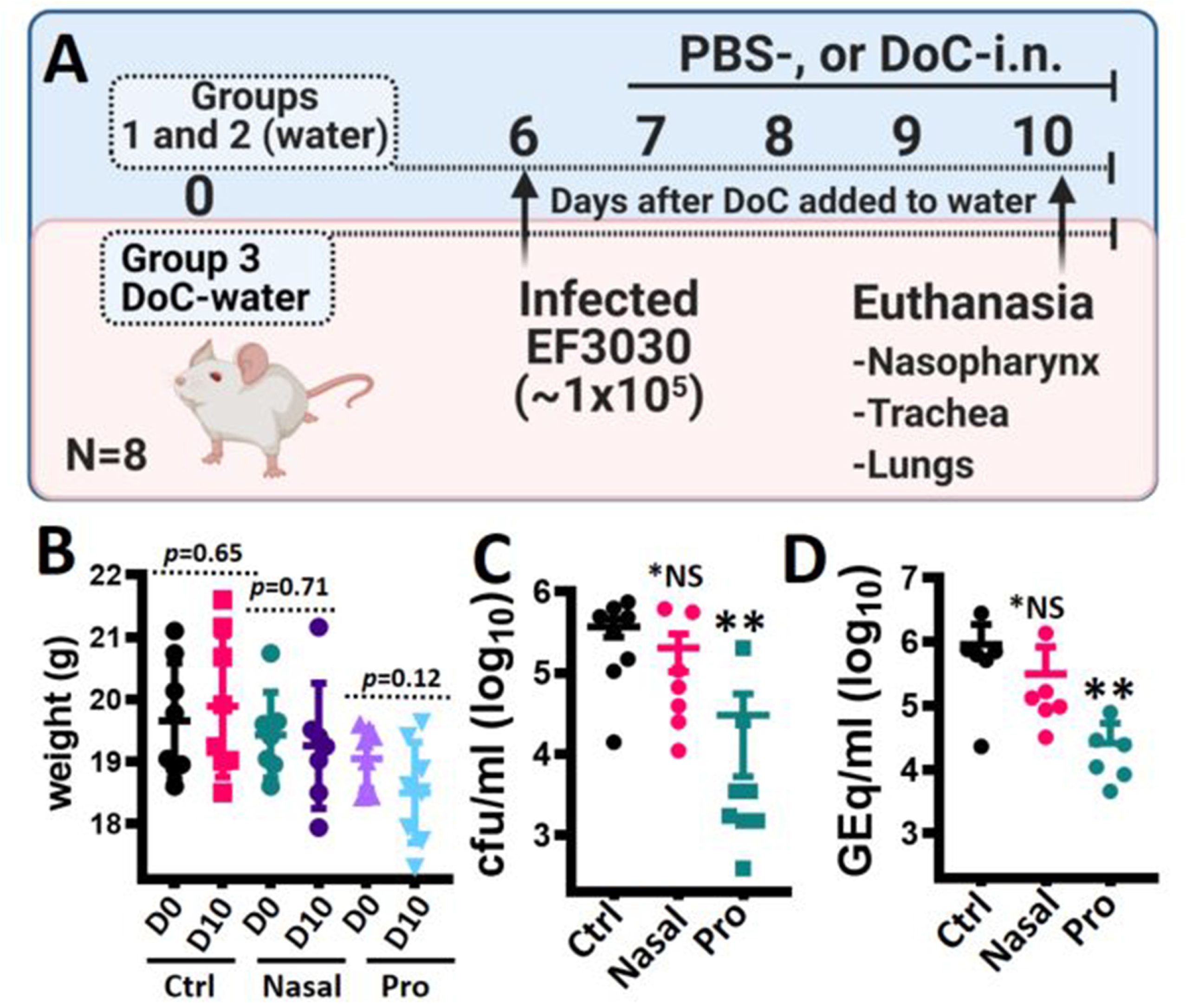
Prophylactic treatment with deoxycholic acid inhibits pneumococcal colonization in a mouse model of colonization. (A) Experimental design. Three groups of mice (N=8) were utilized; groups 1 and 2 drank regular water throughout whereas group 3 was prophylactically (pro) treated by adding DoC (0.2 µg/ml) to their drinking water at day 0. At day 6, all three groups were infected with *S. pneumoniae* EF3030. Twenty-four hours post inoculation, mice in groups 1 and 2 were treated via intranasal (nasal) inoculation with PBS (Ctrl) or DoC (10 µg each nostril) two times a day for four days. All mice were sacrificed at day 10 and the nasopharynx, trachea, lungs and blood were collected. (B) Mice in all three groups were weighed at days 0 and 10. (C) Nasopharyngeal specimens were homogenized, diluted, and plated onto BAP with gentamicin (25 µg/ml) to obtain bacterial density (cfu/ml). (D) DNA was extracted from nasopharyngeal homogenates and used in species-specific *lytA*-based qPCR reactions. In panels B and C =\**p*>0.249 (NS), or \*\**p*<0.003, compared with control mice.

Because of the oral and topical administration of DoC, we monitored the weight of mice daily and no statistically significant difference in weight was observed between day one and the end of the experiment in groups 1, 2, and 3 (Fig. 7B). The colonization density of Spn was determined by dilution and plating of nasopharyngeal homogenates (Fig. 7C). The median density of Spn in the control group was 4.05×10^5^ cfu/ml (25^th^ percentile, 2.50 x10^4^; 75^th^ percentile, 5.60 ×10^5^) whereas mice in the topical DoC-nostril group had a median density of 6.67×10^4^ cfu/ml (25^th^ percentile, 1.15 ×10^5^; 75^th^ percentile, 5.94 ×10^5^). Although slightly reduced compared with the control group, this colonization density was not statistically different. However, mice in the prophylactic DoC administration group had a median Spn density of 2.61 ×10^3^ cfu/ml (25^th^ percentile, 1.5 ×10^3^; 75^th^ percentile, 2.03 ×10^4^) and therefore, a significant reduction of nasopharyngeal colonization density (e.g., 99.36% reduction) was achieved compared to the control group. We further extracted DNA from those nasopharyngeal homogenates and the purified DNA was utilized as template in Spn-specific quantitative (q)PCR reactions. qPCR reactions confirmed a statistically significant decreased density in the oral administration DoC-water group (median, 1.77×10^4^ genome equivalents/ml) compared with the control group (media, 6.70 ×10^5^ genome equivalents/ml) or with the topical DoC-nostril group (media, 1.15 ×10^5^ genome equivalents/ml) (Fig. 7D).

## Discussion

We demonstrated that the prophylactic treatment through the oral route with DoC protected mice from nasopharyngeal colonization with Spn strain EF3030. We also described in the current study a rapid antimicrobial effect of DoC against Spn strains, including reference strains, recent invasive isolates and multidrug resistant strains. DoC-susceptible strains included all PCV13 serotypes, strains bearing resistance to first-line antibiotics utilized to treat pneumococcal disease such as beta-lactams and macrolides (45, 46), and last-resort antibiotics such as meropenem and linezolid. Killing of Spn occurred with 1.27 mM, which is below the upper physiological limit (i.e., 2 mM) of free, unconjugated, bile acids in the intestine although the post-prandial concentration of conjugated bile acids can be as high as 10 mM (47, 48). Remarkably, oral administration of DoC during 10 days by supplementing the drinking water of mice with 0.2 µg/ml (0.5 µM), inhibited nasopharyngeal colonization reducing the pneumococcal density by ∼99%. Since adult mice (20-25 g) drink a minimum of 3 ml of water per day (49), this oral administration via drinking water reached a dosage of ∼0.6 mg/day (∼24 mg/kg/day).

DoC is a secondary bile acid synthesized by the intestinal microbiota from cholic acid and then rapidly absorbed in the intestine (24, 26), thereby it is likely that the concentration of DoC in blood rapidly increased and stayed at similar levels throughout the prophylactic treatment. Whether the level of DoC in circulation directly caused the reduction of the colonization density by means of the DoC-antimicrobial activity, or by means of its immunomodulatory activities, is currently under investigation in our laboratories. Bile produced by mammals has bacteriostatic activity keeping the sterility of the biliary tree, thereby an imbalance in the synthesis of bile acids, among other negative effects, is associated with the overgrowth of bacteria in the small intestine and with inflammation (50). For example, when the intestinal DoC increases the synthesis and secretion of mucus increases and induces the synthesis of immunoregulatory cytokines including the release of human β-defensins (50-52).

DoC is the most abundant secondary bile acid in serum of both mice and humans, with concentrations in healthy subjects ranging from 100 nM to 1 µM (53, 54). Individuals with deficiencies in bile salts have problems to emulsify fat leading to intestinal disorders that have been treated by manipulating intestinal levels of bile acids (51). Cholic acid is used to treat patients with genetic deficiencies in the synthesis of bile acids due to single enzyme deficiencies; the typical dose is 10 to 15 mg/kg once daily (27). Specifically, DoC has been utilized in humans at a concentration of 15 mg/kg/day to decrease plasma high-density lipoprotein (HDL)-cholesterol and low-density lipoprotein (LDL)-cholesterol (27). Thus, the prophylactic dosage of oral DoC that inhibited Spn colonization in mice (24 mg/kg/day) was similar to that utilized to treat metabolic diseases in humans. More recent studies demonstrated that DoC (50-150 µM) and ursodeoxycholic acid (UDA) regulates colonic wound healing using a mouse model of colonic epithelial restitution *in vivo* by administering bile acids at a concentration of 30 mg/kg/day via rectal gavage (55). When administered to mice at a similar concentration, DoC and UDA prevented *C. jejuni*-induced colitis and CDI, respectively. Whereas DoC did not affect viability of *C. jejuni* strains but enhanced an immune response against the pathogen, UDA (3.8 mM) directly decreased the viability of *C. difficile* and directly inhibited sporulation.

Spn strains are “dissolved” in rabbit bile (56), and this is the basis of a phenotypic assay (i.e., bile solubility test) utilized to differentiate Spn strains from other α-hemolytic streptococci (57, 58). At a more physiological concentration, such as that utilized in the current study (1.27 mM), DoC specifically killed Spn strains but had little to no activity against other streptococci. Although the bile solubility test measures turbidity by a subjective visual method rather than viability, bile solubility of streptococci using a semi-quantitative assay correlated with our studies of bacterial viability after a challenge with DoC (59). Similar to our MIC studies, the semi-quantitative assay identified Spn strains having the highest solubility in bile followed by strains with intermediate solubility such as *S. pseudopneumoniae* but all other streptococci were not soluble in bile (59).

Reports describing Spn strains that were not soluble in DoC using the subjective visual readout are available; however, neither the semiquantitative assay nor our viability screening identified strains reduced in or lacking susceptibility to DoC (60). The possibility remains that some pneumococcal strains isolated from pneumococcal disease cases, or those colonizing healthy individuals, are naturally resistant to DoC. Spontaneous resistance to DoC when assessed using strains TIGR4 and EF3030 was not achieved even at bacterial populations >10^12^ cfu/ml. The same strains, when challenged with trimethoprim, generated spontaneous resistant bacteria at a frequency of ≥1.39×10^−9^ A similar spontaneous resistance frequency to trimethoprim (35), to that found in the current study, or spontaneous mutation to optochin (61) have been reported for other Spn strains.

Given the very short treatment with DoC (∼10 min) to reach a MIC *in vitro*, the current study assessed whether topical administration of DoC in the upper airways will result in eradication of colonization. However, mice infected with strain EF3030 and treated with DoC in the nares showed only a slight, but non-significant, reduction of the pneumococcal density. There are a number of reasons to explain the failure to eradicate colonization via the topical route. For example, the small nasal vestibule of mice may have resulted in failure of the DoC to reach all of the nasopharyngeal tissue and/or DoC may have been absorbed before reaching pneumococcal cells. Perhaps a lower pneumococcal carriage density, longer exposure to DoC in the upper airways, or a higher volume of DoC administered into the nostrils would have resulted in a further decrease in bacterial density. Microinvasion of pneumococcus into the nasopharyngeal epithelium observed in this study, and elsewhere (62), could have been also factor for the failure of the topical route.

Earlier biochemical studies suggested that an autolysin(s) was responsible for the lysis of pneumococci in DoC (63). If this is true, such an autolysin should be other than the major LytA or LytC autolysins since our experiments using two different autolysis defective mutants demonstrated similar DoC susceptibility of the R6 wt strain, R6Δ*lytA*Δ*lytC* and R6Δ*spxB*Δ*lctO*. This mechanism, however, appears to be exquisitely specific for Spn strains since challenging other streptococci with the pneumococcus DoC MIC_90_ the majority of those strains were not susceptible. Studies are under way in our laboratories to identify such an enzyme and/or an additional mechanism(s).

In summary, we demonstrated *in vitro* antimicrobial activity of DoC against several pneumococcal strains, including multidrug-resistant strains, *ex vivo* antimicrobial activity to reduce colonization of human nasopharyngeal cells and *in vivo* activity that inhibited colonization in a mouse model of pneumococcal nasopharyngeal colonization. Because Spn strains colonize billions of individuals, killing at least one million every year worldwide, data within this study bears potential for future development of prophylactic interventions aimed to reduce pneumococcal colonization.

## Materials and Methods

### Bacterial strains, culture media and reagents

*Streptococcus* species are listed in Table 1 and Spn reference strains and isogenic mutant derivatives are listed in Table 2. All other Spn are listed in supplemental Table 1. Strains were cultured on blood agar plates containing 5% of sheep red blood cells (BAP) from frozen stocks made in medium containing skim milk, tryptone, glucose, and glycerin (STGG) (64). Animal experiments were cultured on BAP with gentamicin (25 µg/ml). Strains were inoculated in Todd Hewitt broth containing 0.5% (w/v) yeast extract (THY) or in cation-adjusted Mueller-Hinton broth (CAMHB) with 3% of lysed horse blood [(LHB), Remel]. Paraformaldehyde (PFA), gentamicin, tetracycline, trimethoprim, and sodium deoxycholate were sourced from Sigma.

**Table 2.**
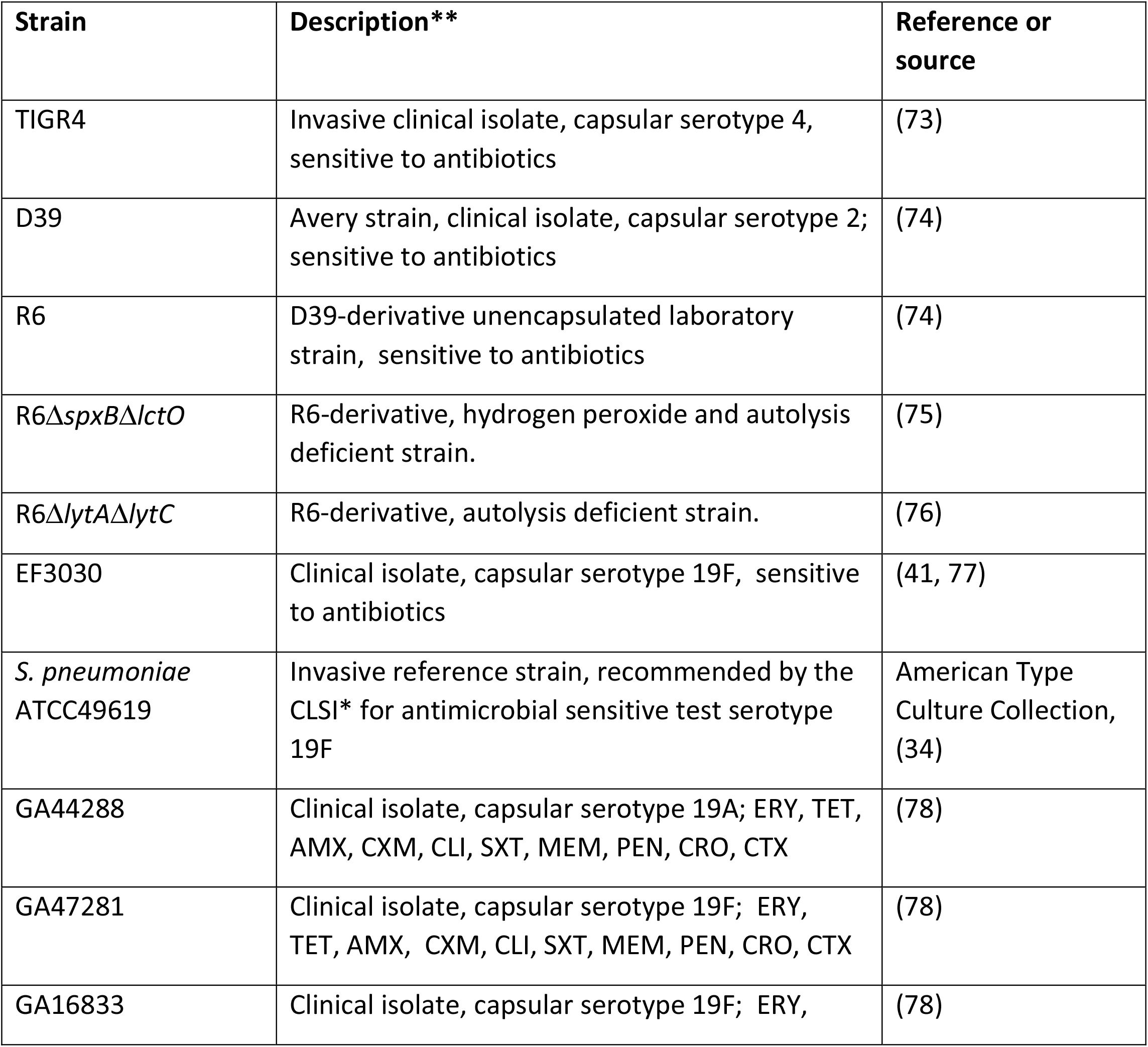
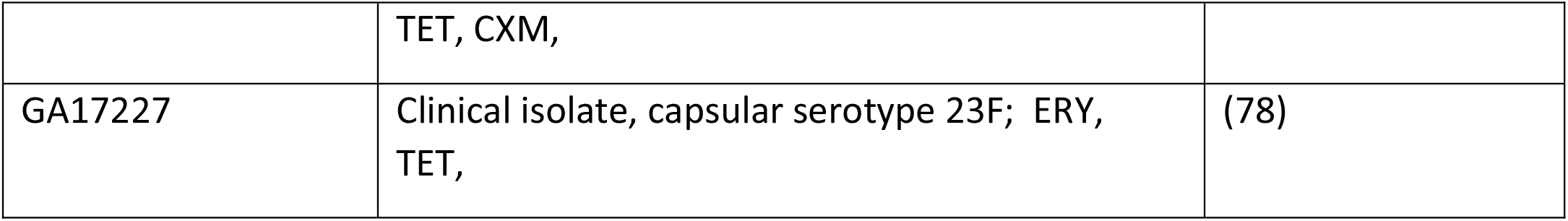
Pneumococcal strains used in this study. laboratory standards institute. **Resistance to: amoxicillin (AMX), cefuroxime (CXM), ceftriaxone (CRO), cefotaxime (CTX), clindamycin (CLI), erythromycin (ERY), meropenem (MEM), penicillin (PEN), tetracycline (TET), trimethoprim-sulfamethoxazole (SXT)

### Preparation of inoculum for experiments

Inoculum was prepared essentially as previously described (65, 66). Briefly, an overnight BAP culture of the strain was used to prepare a bacterial suspension in sterile phosphate buffered saline [(PBS), pH=7.4] and the fresh bacterial suspension was inoculated to a final OD_600_ of ∼0.1. This suspension contained ∼5.15×10^8^ cfu/ml. Aliquots of these suspensions were routinely diluted and plated to confirm bacterial counts (cfu/ml). To inoculate mice, Spn strain EF3030 was inoculated in THY broth and grown until it reached an OD_600_ of ∼0.2 (i.e., early log phase), then sterile glycerol was added to a final concentration of 10% and aliquots were frozen at ∼80°C. An aliquot was removed from each batch to determine the density of the preparations.

### Quantitative studies of the antimicrobial activity of DoC

Studies of antimicrobial activity were performed using THY or CAMHB containing 3% of LHB. Experiments using THY were performed as follows: a bacterial suspension was inoculated in polystyrene 24-well plates (Corning) at a final density of ∼5.15×10^8^ cfu/ml and left untreated (control) or treated with DoC at varying dosages and incubated for 2 h at 37°C in a 5% CO_2_ atmosphere. To remove bacteria that could have potentially attached to the substratum, the microplate was sonicated for 15 s in a Bransonic ultrasonic water bath (Branson, Dunburry CT) and then cultures were serially diluted and plated onto BAP.

To obtain the MIC as recommended by the CLSI we utilized the broth microdilution method (34). DoC was serially diluted in CAMHB containing 3% of LHB in 96-well microtiter plates and pneumococci, that had been adjusted to a turbidity corresponding to the 0.5 McFarland standard (∼1×10^8^ cfu/ml), was inoculated and incubated for 20 h at 37°C. Untreated cultures and non-inoculated medium were included as a control. Besides reading the microplates as recommended by the CLSI, the untreated growth control, and wells with the obtained MIC were serially diluted and plated as before. As a control of the microdilution procedure, the MIC for tetracycline was assessed in parallel using reference strain Spn ATCC49619, and GA16833, which were sensitive (<1 µg/ml) and resistant (8 µg/ml), respectively.

### *Ex-vivo* model of pneumococcal colonization on human pharyngeal cells

This *ex-vivo* adhesion model on immobilized pharyngeal cells was developed by Marks et al. (2012) (67) and thereafter utilized in pathogenesis and biofilm research by different laboratories (67-69). Human pharyngeal Detroit 562 cells (ATCC CCL-198) were cultured in DMEM (Gibco) supplemented with 10% nonheat-inactivated fetal bovine serum (FBS) (Atlanta biologicals), 1% non-essential amino acids (Sigma), 1% glutamine (Sigma), penicillin (100 U/ml), streptomycin (100 µg/ml) and the pH was buffered with HEPES [(10 mM) Gibco]. Cells were incubated at 37°C in a 5% CO_2_ humidified atmosphere until confluence ∼7-10 days on an 8-well glass slide (Lab-Tek), or on CellBIND® surface polystyrene 24-well plates (Corning) and then immobilized by fixation with 2% PFA for 15 min at room temperature. After extensive washes with sterile PBS, immobilized human pharyngeal cells were supplemented with cell culture medium without antibiotics and infected with an inoculum of the tested strain prepared as mentioned earlier. Infected human pharyngeal cells were incubated for 4 h at 37°C and with 5 % CO_2_.

At the end of incubation, planktonic pneumococci were removed, attached bacteria were gently washed two times with sterile PBS and fresh cell culture medium with no antibiotics was added. Pneumococci attached to pharyngeal cells were challenged with DoC at different dosages at incubated for 2 h or treated with 0.5 mg/ml DoC and incubated for the indicated time at 37°C in a 5 % CO_2_ atmosphere. To obtain the density of pneumococci in the 24-well plate model, pneumococci and cells were washed twice with PBS and then sonicated for 15 s in a Bransonic ultrasonic water bath (Branson, Dunburry CT) followed by extensive pipetting to remove attached bacteria. The preparations were diluted and plated onto blood agar plates to obtain bacterial counts (cfu/ml).

To stain pneumococci adhered to cells on the 8-well glass slide, bacteria were fixed with 2% PFA as before and after three washes with PBS the preparations were blocked with 2% bovine serum albumin (BSA) for 1 h at room temperature. These preparations were then incubated for 1 h with serotype-specific polyclonal antibodies (Statens Serum Institute, Denmark) (∼40 μg/ml) that had been previously labeled with Alexa-488 (anti-serotype 4-Alexa-488, to stain TIGR4) or Alexa-555 (anti-serogroup 19-Alexa-555, to stain GA47281) (Molecular Probes). Stained preparations were finally washed two times with PBS. TIGR4 experiments were additionally stained with wheat germ agglutinin conjugated to Alexa-555 [(WGA), 5 µg/ml] and then mounted with ProLong Diamond Antifade mounting medium containing DAPI (Molecular Probes) whereas GA47281 preparations were stained with TO-PRO-3 (1 μM), a carbocyanine monomer nucleic acid stain (Molecular Probes), for 15 min. Confocal images were obtained using a Nikon AX R confocal microscope and analyzed with ImageJ version 1.49k (National Institutes of Health, USA).

### Investigating the spontaneous mutation frequency

To determine the frequency of spontaneous mutation, BAP with 5% sheep red blood cells were prepared to contain either trimethoprim (1 μg/ml) or DoC (0.5 mg/ml). Fresh suspensions of Spn strain EF3030, or TIGR4, made in PBS were prepared at a final density of ∼10^8^, ∼10^9^, ∼10^10^, ∼10^11^, and ∼10^12^ cfu/ml and inoculated on plain BAP or BAP containing trimethoprim or DoC. Bacterial suspensions were diluted and plated onto plain BAP to confirm the density of pneumococci. Inoculated plates were incubated at 37°C under a 5 % CO_2_ atmosphere for ∼20 h. The spontaneous mutation frequency was then calculated by dividing the spontaneous resistant pneumococci, i.e., grown on BAP with trimethoprim or DoC, by the bacterial population.

### Mouse model of pneumococcal nasopharyngeal carriage

Three groups (N=8 each) of inbred 6-7week old C57BL/6 mice (Charles River Laboratories) were utilized to assess *in vivo* antimicrobial activity of DoC. Two groups of mice drank regular water throughout. The drinking water of the third group of mice was supplemented with DoC to a final concentration of 0.2 µg/ml (i.e., 0.02%) starting at day 0 of the experiment, and DoC-containing water was provided *ad libitum* for reminder of the experiment (10 days). Six days after DoC was added to the drinking water of mice in group three, mice in all three groups were anesthetized with 2.5% isoflurane (vol/vol) over oxygen (2 liter/min) administered in a RC2 calibrated vaporizer (VetEquip Incorporated) and then infected with ∼1×10^5^ cfu of Spn EF3030. Twenty-four hours post-nasal-inoculation of EF3030, mice in groups 1 and 2 were treated by nasal instillation with PBS or DoC (10 µg each nostril), respectively, two times a day for four days. Mice were then sacrificed, and the nasopharynx, trachea, lungs and blood were aseptically collected. Tissue homogenates were diluted in PBS and plated onto BAP with gentamicin. Aliquots of these homogenates were supplemented to a final concentration of 10% glycerol and kept at -80°C. The Institutional Animal Care and Use Committee (IACUC) at the University of Mississippi Medical Center approved the protocol used in this study (1584); they oversees the welfare, well-being, and proper care of all mice utilized in this study. All mouse experiments followed the guidelines summarized by the National Science Foundation Animal Welfare Act (AWA).

### DNA extraction from nasopharyngeal homogenates and quantitative (q)PCR reactions

DNA was extracted from mouse nasopharyngeal homogenates using the Qiagen QIAmp Mini Kit. Briefly, an aliquot (50 μl) of nasopharyngeal specimen was added to 100 μL of TE buffer containing 0.04 g/mL lysozyme and 75 U/mL of mutanolysin. Samples were then incubated for 1 h in a 37°C water bath. Following incubation, DNA was extracted from the samples following the recommended protocol from the manufacturer, eluted in 100 μL of buffer AE and kept at -80°C until used. Following extraction, *lyt*A-based qPCR reactions were performed with primers and probe sequences published by the CDC (70), the real-time PCR reagent QuantaBio PerfeCTa FastMix®, and 2.5 μL of DNA template. Reactions were run in duplicate using a CFX96 Real-Time PCR Detection System (Bio-Rad) at the following conditions: 1 cycle at 50°C for 2 min, 1 cycle at 95°C for 2 min and 40 cycles of 95°C for 15 s, and 60°C for 1 min. Standard curves were generated under the same conditions as detailed in our previous studies (71, 72). Considering the genome size of reference strain TIGR4, 2.16 Mb (73), the approximate genome equivalent for each DNA standard was: 4.29×10^5^, 4.29×10^4^, 4.29×10^3^, 4.29×10^2^, 4.29×10^1^, 2.14×10^1^, and 2.14 genome equivalents. Reaction efficiency of standards was within the acceptable range of 90-110%.

### Quantification of extracellular (e)DNA

Spn strains were inoculated into 24-well plates as detailed earlier and incubated for 4 h. The culture supernatants were then harvested by centrifugation for 15 min at 14,000 x *g* in a refrigerated centrifuge (Eppendorf, Hauppauge, NY), and filter sterilized using a syringe-filter (0.4 µm). DNA was purified from 200 ul aliquots of supernatant, as mentioned above, and used as template in *lytA*-based qPCR reactions. For eDNA quantification purposes, standards containing 1×10^3^, 1×10^2^, 1×10^1^, 1×10^0^, 1×10^−1^, 5×10^−2^, 1×10^−3^ pg of chromosomal DNA purified from strain TIGR4 were run in parallel to generate a standard curve. The standard curve, and regression equation obtained, was then used to calculate final pg/ml using the CFX software (Bio-Rad, Hercules, CA).

### Statistical analysis

Statistical analysis was performed by the non-parametric two-tailed Student *t* test, or the Mann-Whitney *U* test (comparing two groups) using the software GraphPad Prism version 9.0.0 (121).

## Figure legends

**Supplemental figure 1. Deoxycholic acid kill *S. pneumoniae* strain TIGR4 inoculated into CAMHB with 3% lysed horse blood.** *S. pneumoniae* strain TIGR4 was inoculated at a density of ∼5.15×10^8^ cfu/ml into CAMHB-LHB and left untreated (control) or treated with 0.5, 1, or 2 mg/ml of deoxycholic acid (DoC). Bacteria were incubated ∼20 h at 37°C in a 5% CO_2_ atmosphere after which the cultures were serially diluted and plated onto blood agar plates to obtain the density (cfu/ml). Error bars represent the standard errors of the means calculated using data from at least three independent experiments. The median density is shown inside the control bar or above DoC (0.5). Limit of detection (LOD) was <10 cfu/ml.

## Acknowledgements

This study was in part supported by grants from the National Institutes of Health (NIH; 1R21AI144571-01, and 1R21AI151571-01A1 to JEV). BA was supported by a Fulbright scholarship awarded by the US Department of State. The content is solely the responsibility of the authors and does not necessarily represent the official view of the NIH or the US Department of State. We thank Dr. Yih-Ling Tzeng and Dr. David Stephens from Emory University School of Medicine and Dr. Lesley McGee from the Centers for Disease Control and Prevention (CDC) for providing *S. pneumoniae* strains. We also thanks Dr. Miriam Moscoso from “Centro de Investigaciones Biologicas” in Madrid, Spain for providing the autolysin knockout mutant. Special thanks to Dr. Gene L. Bidwell (Cell and Molecular Biology) and Joshua R. Jefferson (Pathology) at UMMC for providing assistance with confocal microscopy and histology studies, respectively. Landon Murin from the Murrah-UMMC base pair program for his assistance in some laboratory procedures.

